# Crop yields under simulated nuclear winter: a growth chamber experiment

**DOI:** 10.64898/2026.05.05.723012

**Authors:** Simon Blouin, Desa Rae Abrams, Rotem Ben-Zeev, Charles T. Anderson, Jesse R. Lasky, David Denkenberger

## Abstract

A global nuclear war could inject soot into the stratosphere, blocking sunlight and causing rapid cooling. Assessments of the resulting agricultural collapse rely on crop models never validated under such conditions. We grew wheat, canola, and potato in growth chambers simulating the light and temperature of an extreme nuclear winter at tropical and temperate sites. In the tropical chamber (18–20 °C, 200 μmol m^-2^ s^-1^ PAR), all three crops produced viable yields. Wheat yielded 2.1–2.3 t/ha (*n*=3 well-watered, *n*=3 water-stressed pots), 60% of the global average, and single-pot observations of canola and potato suggested biological yields comparable to global averages. In the temperate chamber simulating nuclear winter irradiance (60–360 μmol m^−2^ s^−1^), wheat stems collapsed under their own weight. Although hand-harvesting recovered 0.6–2.8 t/ha of grain, mechanical field harvest of a flat canopy would recover substantially less. This failure mode was not observed in a higher-light control chamber and is not captured by existing crop models, which may therefore overestimate temperate cereal production under nuclear winter. Canola produced comparable yields under both temperate light regimes without lodging. Empirical screening of additional staples is needed to identify which remain viable under nuclear winter.

## Introduction

A nuclear war between major powers is among the most serious threats to global food security (Xia *et al*., 2022). The direct effects (blast, fire, and radioactive fallout) would be devastating but geographically confined (Robock *et al*., 2023). The largest threat to human civilization may be the resulting climate disruption. Burning cities could inject soot into the stratosphere, which would block sunlight and cause rapid global cooling, a phenomenon known as nuclear winter (Robock *et al*., 2007; Coupe *et al*., 2019). Climate simulations of a 150 Tg soot injection scenario, corresponding to a worst-case all-out war between major nuclear powers, predict that global-average cropland temperatures would fall by approximately 15 °C and solar radiation by approximately 80% during the worst years, with effects persisting for more than a decade (Coupe *et al*., 2019; Xia *et al*., 2022). Crop modelling driven by these climate projections suggests that global caloric production from crops could diminish by approximately 90% within a few years after the war, placing billions of people at risk of starvation (Xia *et al*., 2022). Without adaptation, global caloric production would fall below the minimum required to feed the world population for six consecutive years under the 150 Tg scenario (Blouin *et al*., 2025). A recent review commissioned by the U.S. Congress concluded that the potential sun-obscuring effects of nuclear war remain a serious concern warranting continued research (National Academies of Sciences, Engineering, and Medicine, 2025).

Nuclear winter yield projections rely on process-based crop models. For instance, Xia *et al*. (2022) used CLM5crop (Lombardozzi *et al*., 2020), Blouin *et al*. (2025) used DSSAT (Jones *et al*., 2003), and Shi *et al*. (2025) used Cycles (Kemanian *et al*. 2024) to simulate major crops under a 150 Tg nuclear winter. However, these models were calibrated under current-climate conditions and have never been validated at nuclear-winter-relevant extremes. Crop models lose accuracy when extrapolated beyond their calibration envelope. For example, prediction failures during moderate extremes, such as the 2003 European heatwave (Schewe *et al*., 2019), demonstrate that models can fail even within the range of historical climate variability.

Forty years of nuclear winter research has produced no published empirical yield data for staple crops under the relevant environmental conditions. Femeena *et al*. (2023) demonstrated that duckweed (family *Lemnaceae*) can accumulate biomass under extreme low light (as low as 7 μmol m^-2^ s^-1^) and proposed it as a protein source for post-catastrophic food production, but duckweed is not representative of the staple crops that currently underpin global calorie supply. A substantial literature on partial-season shading of grain crops exists, with yield reductions of 30 to 70% reported under 50 to 75% light reduction applied during individual growth phases (Fischer, 1975; Willey and Holliday, 1971; McMaster *et al*., 1987; Savin and Slafer, 1991; Dong *et al*. 2014; Artru *et al*., 2017; Islam *et al*., 2021; Jia *et al*., 2021). However, these studies applied low-light conditions over periods of weeks to months; none subjected crops to a full growing season under sustained low light. We are not aware of published experiments that have grown any staple crop under combined nuclear winter conditions (low light and cold temperatures) for a full growing season.

The objective of the present study is to provide the first empirical yield data for staple crops under simulated nuclear winter conditions, by growing wheat, canola, and potato in controlled environment chambers reproducing the low light and cold temperatures predicted for tropical and temperate sites. These data provide an empirical baseline for the crop models that underpin published assessments of food security under nuclear winter conditions, and can inform adaptation planning, including the selection of viable crops and cultivars for regions that maintain agricultural viability under nuclear winter (Blouin *et al*., 2025; McLaughlin *et al*., 2025; Shi *et al*., 2025).

## Materials and methods

### Experimental design

Three growth chambers at the Pennsylvania State University, USA were used to grow wheat (*Triticum aestivum* L.), canola (*Brassica napus* L.), and potato (*Solanum tuberosum* L.) under conditions simulating nuclear winter from May 2025 to March 2026. One Conviron walk-in growth chamber equipped with a CMP4030 control system (Controlled Environments Ltd, Winnipeg, MB, Canada; hereafter, the “tropical chamber”) simulated nuclear winter conditions in the tropics (Supplementary Fig. 1A). Two Percival reach-in growth chambers (model AR66L3; Percival Scientific, Perry, IA, USA; hereafter, the “temperate chambers”) simulated nuclear winter conditions at a temperate latitude; both shared the same temperature regime but one received the predicted nuclear winter irradiance (low-light) and the other a higher-light control, isolating the effect of reduced radiation (Supplementary Fig. 1B). Within each chamber, pots were assigned to either a well-watered or a water-stressed treatment.

The temperate scenario simulated conditions at a site in South Australia during the worst year of a 150 Tg nuclear winter, based on climate projections from Coupe *et al*. (2019) (including a debiasing correction as in Blouin *et al*., 2025). Crop modelling of the 150 Tg scenario predicts that Australia, along with other Southern Hemisphere nations, retains significant agricultural capacity under nuclear winter (Xia *et al*., 2022; Blouin *et al*., 2025). However, these predictions rely on models that have never been validated under such extreme conditions, motivating empirical testing at a representative temperate Southern Hemisphere site.

The two temperate chambers reproduced a month-by-month progression of temperature and photoperiod (Fig. 1), including a five-week vernalization period (27 June to 8 August 2025; 4 °C, 60 μmol m^-2^ s^-1^ PAR, 10 h photoperiod), satisfying the requirements of both winter crops. The lower limit for chamber temperatures was set at 4 °C. Although the climate projections predict lower temperatures during the coldest months, the objective of this experiment was to quantify yield under growing season conditions predicted to occur during a nuclear winter, not to test overwintering survival. Crop growth effectively ceases around or below 4 °C (Salazar-Gutierrez *et al*., 2013), and the winter-hardy cultivars used (TAM 114, Wichita) have been released for commercial production in the U.S. Great Plains where they are exposed to sub-zero temperatures. The low-light chamber received the light intensities predicted during nuclear winter (60 to 360 μmol m^-2^ s^-1^ depending on month), while the high-light chamber received a constant 360 μmol m^-2^ s^-1^ as a control. The maximum PAR output of the Percival chambers was 360 μmol m^-2^ s^-1^; nuclear winter predictions exceeded this value during the final months of the growing season, so during those months both chambers received the same irradiance. CO_2_ concentration was not actively controlled in the sealed Percival chambers and fluctuated over the course of the experiment, averaging 552 μmol/mol.

**Fig 1.**
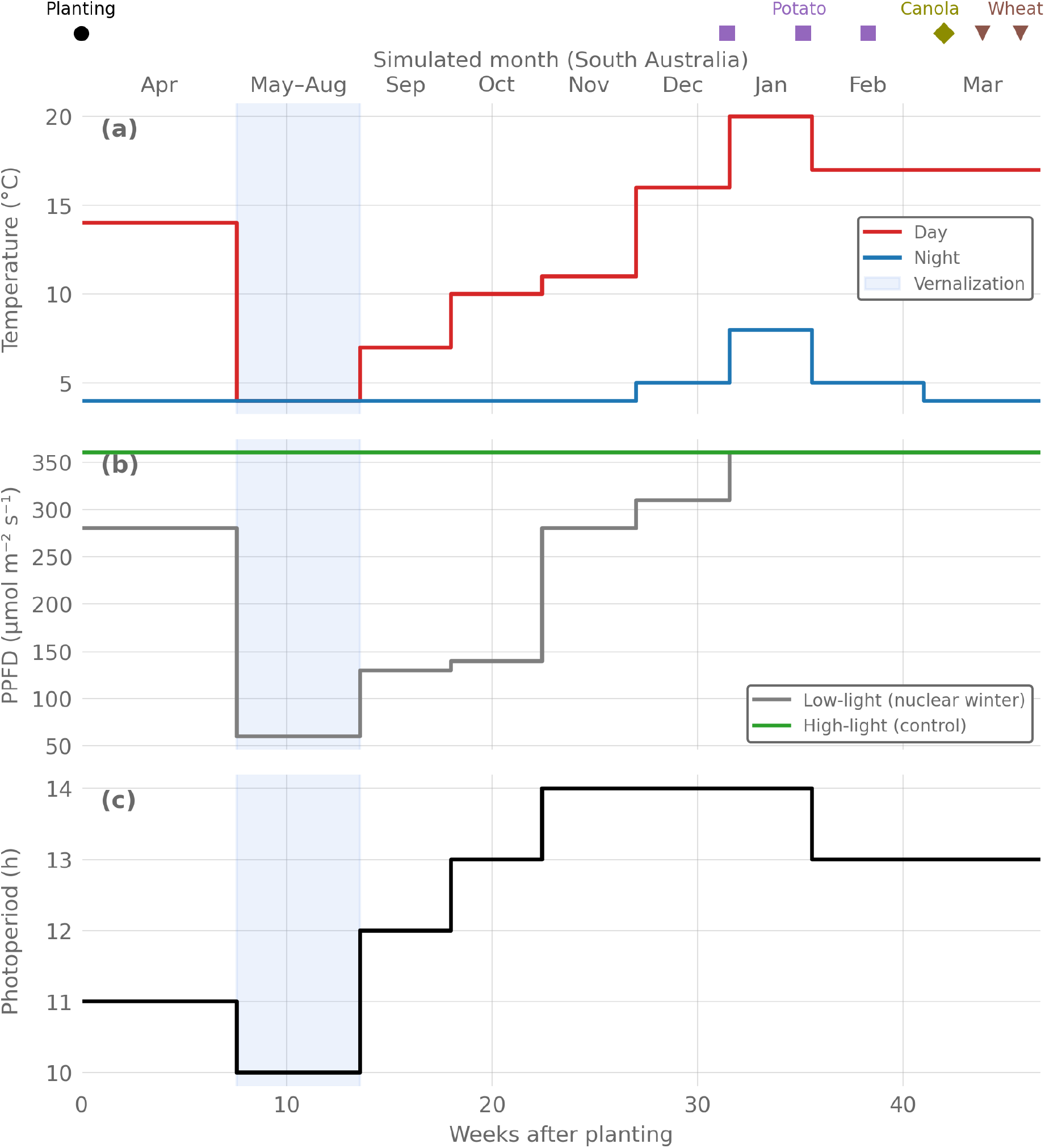
Environmental conditions in the temperate chambers over the experimental period. (*A*) Day and night temperature setpoints. (*B*) PAR for the low-light (nuclear winter) and high-light (control) chambers. (*C*) Photoperiod (identical in both chambers). The blue shading indicates the vernalization period. Simulated Southern Hemisphere months (South Australia) are shown along the top axis. Markers indicate planting (circle) and harvest dates for potato (squares), canola (diamond), and wheat (triangles); where multiple markers of the same shape appear, they correspond to separate harvest dates across chambers or water treatments. The tropical chamber (not shown) maintained constant conditions throughout: 18–20 °C, 200 μmol m^-2^ s^-1^ PAR, 16 h photoperiod.

The tropical chamber simulated equatorial conditions (proxy for Indonesia). Tropical regions are similarly predicted to maintain meaningful crop yields under nuclear winter (Xia *et al*., 2022; Blouin *et al*., 2025). The chamber was held at constant 18 to 20 °C with 200 μmol m^-2^ s^-1^ PAR and a 16 h photoperiod. The 16 h photoperiod at 200 μmol m^-2^ s^-1^ provided a daily light integral equivalent to 12 h at approximately 267 μmol m^-2^ s^-1^, comparable to the predicted equatorial irradiance during the worst year of the 150 Tg scenario. The longer photoperiod was dictated by shared use of the chamber.

Within each chamber, pots were assigned to one of two water treatments: well-watered, irrigated to approximately 70% plant-available water (PAW); or water-stressed, maintained at approximately 40% PAW. The water-stressed target was based on crop model predictions for rainfed conditions at the temperate site during the worst year of a 150 Tg scenario (Blouin *et al*., 2025). Post-catastrophic water availability is highly uncertain, depending on regional precipitation changes, local hydrology, and whether irrigation infrastructure survives the conflict. Including a water-stressed treatment alongside the well-watered control allowed the experiment to probe whether water limitation compounds the effects of cold and low irradiance, a question that cannot be resolved from single-stress experiments because plant responses to combined abiotic stresses are not additive (Mittler, 2006). The 40% PAW target sits just below the soil water threshold at which crop transpiration and yield begin to decline for cereals (Allen *et al*., 1998), making it a representative moderate stress without imposing near-wilting conditions.

### Plant material

Cultivars were selected for early maturity and adaptation to cold climates. The temperate chambers grew winter wheat cv. TAM 114, a hard red wheat with good winter hardiness (Rudd *et al*., 2018); winter canola cv. Wichita, a winter-hardy canola developed for the U.S. Great Plains (Rife *et al*., 2001); and potato cv. Red Norland, an early-maturing cultivar grown in cold-climate regions (Johansen *et al*., 1959). The tropical chamber grew spring wheat cv. Glenn, a medium-early hard red wheat (Mergoum *et al*., 2006); the ultra-early spring canola hybrid NC155 TF (Nuseed; Fordyce *et al*., 2023); and potato cv. Red Norland. Seeds and tubers were planted on 5 May 2025.

Each pot (30.5 × 30.5 × 30.5 cm) contained nine wheat plants (corresponding to approximately 100 plants/m^2^, within the range of recommended field densities; Whaley *et al*., 2000), six canola plants (approximately 65 plants/m^2^, typical of commercial target densities; Hartman and Jeffrey, 2021), or one potato plant (approximately 11 plants/m^2^, roughly twice the typical field density due to single-pot constraints; Hou *et al*., 2020). Each temperate chamber held ten pots and the tropical chamber held eleven. Wheat was replicated with three pots per treatment per chamber; canola and potato had one pot per treatment per chamber, except that the tropical chamber had two water-stressed canola pots and one well-watered canola pot. Canola and potato results are reported as exploratory. The temperate temperature regime included prolonged cold periods designed for winter crops; potato plants were included in these chambers to observe their response to a temperate nuclear winter scenario, despite the conditions being poorly suited to potato production.

### Growing medium, data collection, and harvest

The growing medium was a 1:1:1 mixture (by volume) of PRO-MIX BX potting mix, 20–40 mesh silica sand, and Turface calcined clay. Osmocote slow-release fertilizer was incorporated into the growing medium. The water-stressed treatment was initiated after germination to ensure seedling establishment. Plant height was measured weekly on three tagged plants per pot for wheat and canola and all plants for potato.

Maturity was assessed by weekly visual inspections. Wheat pots were harvested when 90% of heads had reached the straw-ripe stage (yellow heads, grains hard). Canola pots were harvested when 90% of pods had turned brown, except for the tropical spring canola (see below). Potato was harvested when the foliage had fully senesced. At maturity, wheat heads were clipped, threshed, and grain weighed. A 100-grain subsample was dried at 105 °C for 24 h to determine moisture content; grain yields are reported at 135 g/kg moisture. Canola pods were clipped, threshed, and seeds weighed; a 500-seed subsample was dried to determine moisture content; seed yields are reported at 90 g/kg moisture. Potato tubers were hand-collected and weighed; potato yields are reported as fresh weight.

Spring canola in the tropical chamber exhibited indeterminate flowering under the favourable constant conditions, with plants continuing to initiate new pods while earlier pods matured. Because dried pods shattered and dispersed seeds before the remaining pods ripened, mature pods were clipped progressively into labelled bags as they dried. Once approximately 50% of pods had reached maturity, whole canopies were cut and cured in paper bags for 7 to 10 days before threshing, analogous to swathing in field practice.

## Statistical analysis

Wheat was replicated with three pots per water treatment per chamber, and treatment means are reported with standard errors. Differences between water and light treatments were tested using Welch’s *t*-test. Canola and potato had one pot per treatment per chamber and are reported as single-pot observations without statistical testing.

## Results

### Growth and phenology

All crops germinated successfully in all three chambers. Seedlings emerged within 7 days of planting in the tropical chamber and within 18 days in the temperate chambers, where initial temperatures were lower. All pots were thinned to the target densities described in Materials and methods.

In the tropical chamber, spring wheat reached a peak mean height of 76 ± 2 cm (mean ± s.e., *n* = 6 pots) by week 13 and was harvest-ready by week 21. Spring canola flowered continuously from week 8 onwards. Potato plants flowered by week 6 and foliage had senesced by week 22.

In the temperate chambers, growth was slow during the cold months and the five-week vernalization period, then accelerated once temperatures rose above 10 °C (simulated October onwards; Fig. 1). Extended height trajectories for the winter wheat in the two temperate chambers are shown in Fig. 2. During the first months, low-light wheat plants were taller than high-light plants. This trend reversed from approximately week 22 onwards: high-light, well-watered wheat reached 118 ± 2 cm and produced 63 ± 4 heads per pot, while high-light, water-stressed wheat reached 104 ± 10 cm with 51 ± 4 heads per pot. Low-light wheat plateaued at 93 ± 13 cm (well-watered) and 86 ± 5 cm (water-stressed), with 42 ± 2 and 18 ± 6 heads per pot, respectively. All values are means ± s.e. (*n* = 3 pots). From approximately week 19 onwards, well-watered wheat plants were taller than water-stressed plants in both chambers.

**Fig 2.**
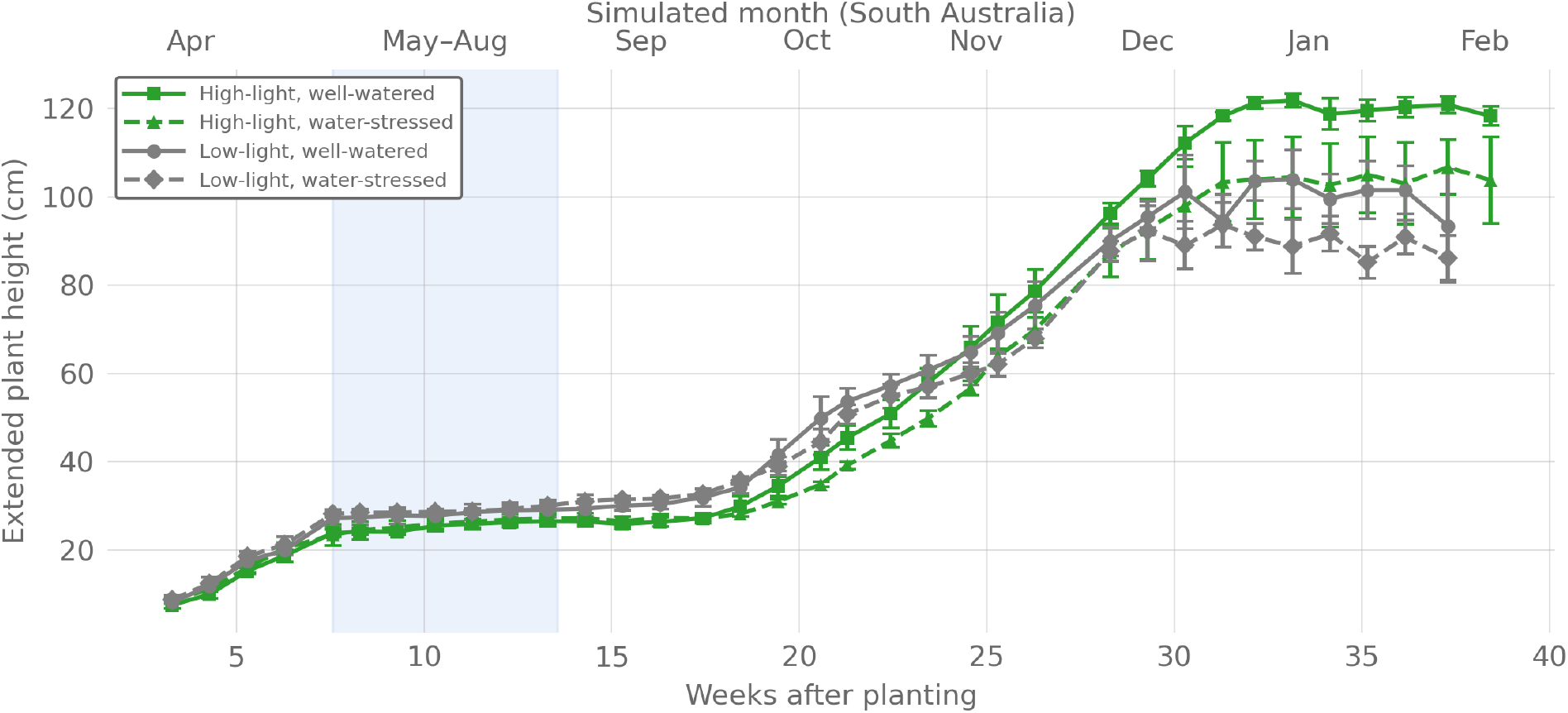
Extended plant height over time for winter wheat (cv. TAM 114) in the two temperate chambers. Extended plant height is measured from base to tip along the stem. The high-light chamber (green) received a constant 360 μmol m^-2^ s^-1^ PAR; the low-light chamber (grey) received the nuclear winter irradiance shown in Fig. 1*B* (60–360 μmol m^-2^ s^-1^ depending on month). Solid lines denote the well-watered treatment; dashed lines denote the water-stressed treatment. Values are means with s.e. bars (*n* = 3 pots, each the mean of three tagged plants). The blue shading indicates the vernalization period. Extended plant height rather than vertical height is reported because low-light plants progressively lost the ability to remain upright (see Fig. 3).

Winter canola began flowering by week 28 in both temperate chambers and set pods normally. Well-watered potato foliage senesced and was harvested between weeks 35 and 38. Water-stressed potato plants in both temperate chambers died rapidly from weeks 28 to 29 following an irrigation event. The symptoms of this die-off are consistent with vascular wilt. These plants were harvested at week 31.

Smut infection was observed in tropical spring wheat, affecting approximately 20% of heads in one well-watered pot and approximately 5% in one water-stressed pot; remaining pots were unaffected. Thrips were detected in the tropical chamber at week 10 and treated with spinosad (Conserve SC) and predatory mites (*Amblyseius cucumeris*). Fungus gnats were also detected in the tropical chamber and treated with entomopathogenic nematodes. No pest or disease issues were observed in the temperate chambers apart from the potato vascular wilt described above.

### Stem integrity in temperate low-light wheat

From week 26 onwards, wheat stems in the low-light temperate chamber began bending at the nodes and collapsing under their own weight (Fig. 3*B*). The collapse was progressive, and by week 38 the canopy was completely matted. No stem collapse was observed in the high-light chamber under otherwise identical conditions (Fig. 3*A*). The growth chambers were sealed environments with no wind, rain, or mechanical disturbance; the structural failure occurred under the plant’s own weight. Up to the point of collapse, the low-light chamber had received 60 to 280 μmol m^-2^ s^-1^ PAR (Fig. 1*B*). Stem collapse began before the low-light schedule reached the 360 μmol m^-2^ s^-1^ chamber ceiling (see Materials and methods), so the failure occurred at the irradiance levels predicted by the climate model for this site, not at artificially reduced values. By contrast, the high-light chamber received 360 μmol m^-2^ s^-1^ throughout.

**Fig 3.**
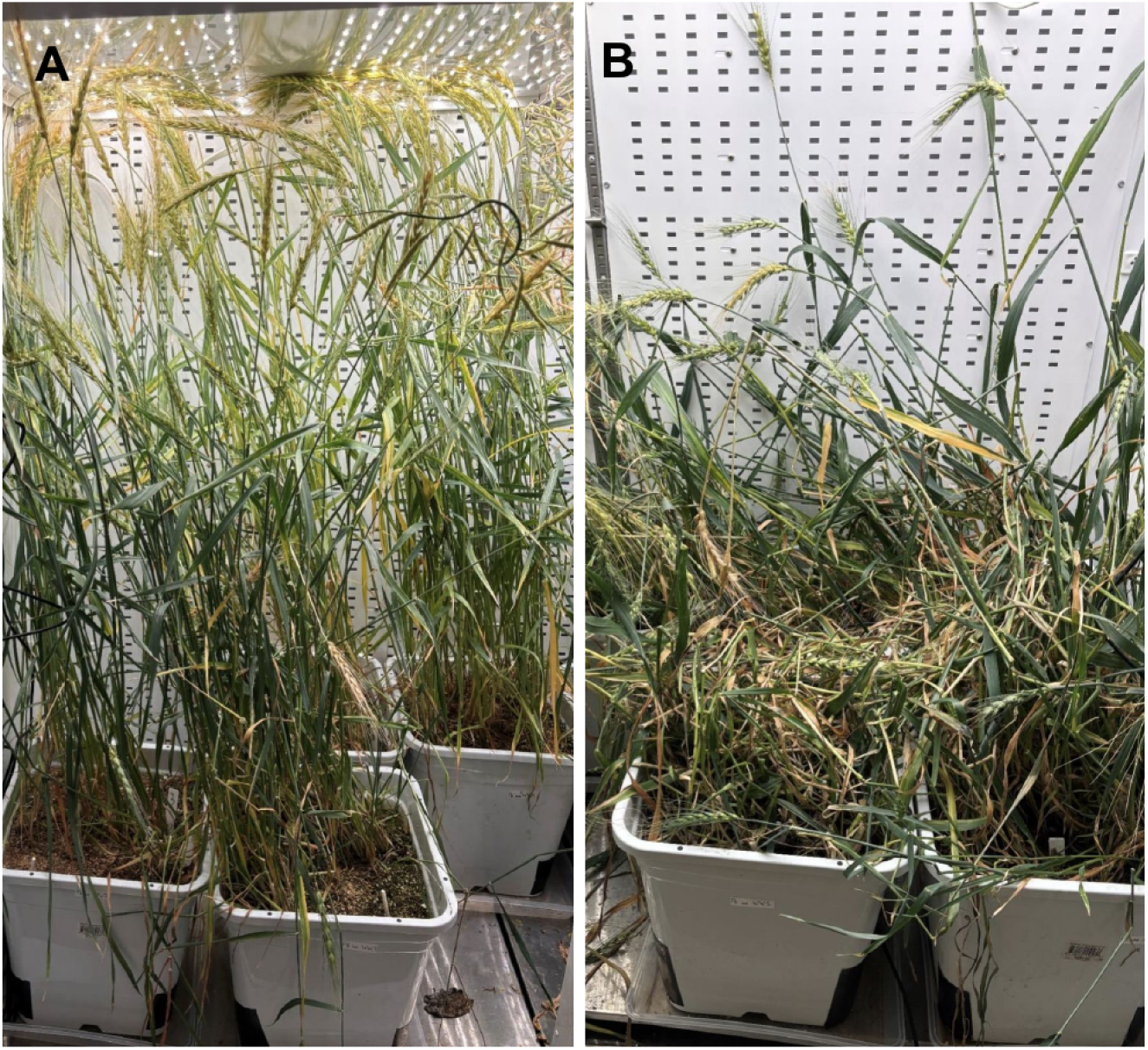
Winter wheat (cv. TAM 114) in the temperate chambers 37 weeks after planting. (*A*) High-light control chamber (360 μmol m^-2^ s^-1^ PAR): stems remained upright with well-formed heads. (*B*) Low-light chamber (nuclear winter irradiance, 60–360 μmol m^-2^ s^-1^ PAR depending on month): stems collapsed under their own weight.

### Harvest yields

Harvest yields for all crops, chambers, and water treatments are presented in Table 1. Yields are expressed as field-scale equivalents based on pot surface area (0.093 m^2^). They should not be directly interpreted as achievable field yields, because pot-based experiments are subject to root restriction artifacts (Poorter *et al*., 2012), elevated CO_2_, and lack the wind, rain, weed, and pest pressures of field conditions (Passioura, 2006).

**Table 1.** Harvest yields for crops grown under simulated nuclear winter conditions in the three growth chambers described in Materials and methods. Adjusted mass is the grain, seed, or tuber mass per pot after moisture standardization (wheat at 135 g/kg, canola at 90 g/kg, potato as fresh weight). Yield is the field-scale equivalent calculated from the pot area (0.093 m^2^). Wheat values are means with s.e. (*n* = 3 pots per treatment). All other entries are single-pot observations (*n* = 1 unless noted).

| Crop | Adjusted mass,<br>well-watered<br>(g/pot) | Yield,<br>well-watered<br>(t/ha) | Adjusted mass,<br>water-stressed<br>(g/pot) | Yield,<br>water-stressed<br>(t/ha) |
| --- | --- | --- | --- | --- |
| ***Tropical chamber*** |  |  |  |  |
| Spring wheat (Glenn) | 19.0 (0.9) | 2.05 (0.10) | 21.6 (0.2) | 2.32 (0.03) |
| Spring canola (NC155 TF) | 18.9 | 2.03 | 20.3<sup>a</sup> | 2.19<sup>a</sup> |
| Potato (Red Norland) | 516 | 55.5 | 218 | 23.5 |
| ***Temperate high-light chamber*** |  |  |  |  |
| Winter wheat (TAM 114) | 12.6 (5.5) | 1.35 (0.59) | 8.0 (3.5) | 0.86 (0.37) |
| Winter canola (Wichita) | 28.1 | 3.03 | 24.6 | 2.64 |
| Potato (Red Norland) | 214 | 23.0 | 28.2<sup>b</sup> | 3.0<sup>b</sup> |
| ***Temperate low-light chamber*** |  |  |  |  |
| Winter wheat (TAM 114) | 26.1 (4.1) | 2.81 (0.44) | 5.8 (3.7) | 0.62 (0.40) |
| Winter canola (Wichita) | 29.0 | 3.11 | 20.4 | 2.19 |
| Potato (Red Norland) | 138 | 14.8 | 102<sup>b</sup> | 11.0<sup>b</sup> |

<sup>a</sup> Mean of two water-stressed pots ( $n = 2$ ). Individual pot yields were 2.28 and 2.09 t/ha.
<sup>b</sup> Water-stressed potato plants in the temperate chambers were compromised by vascular wilt.

In the tropical chamber, spring wheat yielded 2.05 ± 0.10 t/ha (well-watered) and 2.32 ± 0.03 t/ha (water-stressed). This difference was not statistically significant (*p* = 0.10, *t*-test). In single-pot observations, spring canola yielded approximately 2 t/ha in both water treatments, and potato yielded 55.5 t/ha (well-watered) and 23.5 t/ha (water-stressed).

In the temperate high-light chamber, winter wheat yielded 1.35 ± 0.59 t/ha (well-watered) and 0.86 ± 0.37 t/ha (water-stressed), with large pot-to-pot variability. This difference was not statistically significant (*p* = 0.53, *t*-test). Many heads were partially or entirely empty, contributing to large pot-to-pot variability in grain yield despite more consistent head counts. In single-pot observations, winter canola yielded 3.03 t/ha (well-watered) and 2.64 t/ha (water-stressed). Well-watered potato yielded 23.0 t/ha; the water-stressed potato yield of 3.0 t/ha was compromised by vascular wilt (both single-pot observations).

In the temperate low-light chamber, winter canola yielded 3.11 t/ha (well-watered) and 2.19 t/ha (water-stressed) in single-pot observations. Potato yielded 14.8 and 11.0 t/ha (single-pot observations), though the water-stressed value was again compromised by vascular wilt. Winter wheat in the low-light chamber yielded 2.81 ± 0.44 t/ha (well-watered) and 0.62 ± 0.40 t/ha (water-stressed); the difference between water treatments was statistically significant (*p* = 0.02). The water-stressed treatment showed high pot-to-pot variability, with individual pots yielding 0.07, 0.41, and 1.39 t/ha. These yields were obtained by hand-harvesting individual heads from the matted canopy (see Stem integrity in temperate low-light wheat, above). They represent biological grain production, and mechanical harvest of severely lodged canopies would recover substantially less grain (Berry *et al*., 2004).

## Discussion

### Tropical yields

The tropical chamber results suggest that staple crops can produce meaningful yields under the light and temperature conditions predicted for the worst year of a 150 Tg nuclear winter at equatorial latitudes. Spring wheat yielded approximately 2 t/ha, comparable to global average wheat yields in the 1980s and roughly 60% of the current global average of 3.6 t/ha (Ritchie *et al*., 2023). Tropical canola yielded approximately 2 t/ha, near the current global average of 2.1 t/ha, and tropical potato yielded 23.5 to 55.5 t/ha depending on water treatment, in the range of the current global average of 22.9 t/ha (Ritchie *et al*., 2023). The canola and potato values are single-pot observations, but they suggest that economically viable yields are plausible for these crops under simulated nuclear winter conditions.

These results are broadly consistent with crop model predictions that tropical regions retain significant agricultural capacity under a 150 Tg nuclear winter (Xia *et al*., 2022; Blouin *et al*., 2025). However, these yields should be interpreted as demonstrating biological feasibility rather than as field yield predictions. The growth chambers lacked wind, weeds, and natural pest pressure, all of which would reduce yields under real post-war conditions.

### Wheat stem failure under low light

A key finding of this experiment is the structural failure of wheat stems (lodging) under temperate low-light conditions. Under nuclear winter irradiance, wheat stems bent at the nodes and collapsed under their own weight without any wind, rain, or mechanical disturbance (Fig. 3*B*). The collapse was observed only in the low-light chamber; the high-light control chamber, which shared the same temperature regime, produced upright plants (Fig. 3*A*), demonstrating that cold temperatures alone did not cause stem failure.

Hand-harvesting individual heads from the matted low-light canopy recovered 2.81 ± 0.44 t/ha (well-watered) and 0.62 ± 0.40 t/ha (water-stressed) of grain (Table 1). The high-light control produced 1.35 ± 0.59 t/ha (well-watered) and 0.86 ± 0.37 t/ha (water-stressed). The nominally higher well-watered yield in the low-light chamber does not indicate a benefit of reduced irradiance. The difference between the low- and high-light well-watered yields was not statistically significant (*p* = 0.12), the high-light treatment exhibited large pot-to-pot variability, and, most importantly, the low-light grain was recoverable only because each head was collected individually by hand. Mechanical harvest of severely lodged wheat suffers large losses because the combine header cannot engage a flat canopy (Berry *et al*., 2004), so effective field yield under nuclear winter irradiance would be far below the hand-harvested values reported here.

Lodging is conventionally attributed to external forces such as wind (Pinthus, 1974). In this case, stems failed under their own weight, in a sealed chamber with minimal air movement, indicating that the low-light environment weakened stems to the point where gravitational loading alone was sufficient to cause collapse. The underlying mechanism was not directly measured in this experiment, but is well supported by the existing literature. For instance, Luo *et al*. (2022) found that 50% shading with black nylon netting (reducing PAR from approximately 1100 to 550 μmol m^-2^ s^-1^) decreased lignin content of wheat stems by 14 to 29% (depending on shading duration and planting density), thereby decreasing lodging resistance since lignin content is strongly correlated with stem breaking strength (Zheng *et al*., 2017). Our low-light chamber provided 60 to 280 μmol m^-2^ s^-1^ up to the point of stem collapse — far more severe than the 50% shading applied by Luo *et al*. (2022) — and would be expected to produce correspondingly greater lignin depletion and structural weakness. Shading also promotes stem elongation (Luo *et al*., 2022), compounding the structural problem by increasing the mechanical lever arm. This pattern was observed in the present experiment: low-light plants were initially taller than high-light plants (Fig. 2) before their stems began to fail.

Because the growth chambers lacked wind and rain, field-grown wheat would experience additional mechanical loading that would compound the risk of lodging. However, the absence of wind also means that our chamber-grown plants lacked thigmomorphogenetic reinforcement. Hindhaugh *et al*. (2021) showed that mechanical stimulation nearly doubles the Young’s modulus of wheat stem segments. But it is unlikely that this approximate doubling in stem stiffness compensates for the addition of wind loading under field conditions (wind is normally the dominant cause of cereal lodging; Baker, 1995). Winter canola in the low-light chamber did not exhibit stem failure and produced yields comparable to the high-light control in single-pot observations (3.11 vs 3.03 t/ha, well-watered; Table 1), raising the possibility that its branched architecture confers greater structural resilience under low light.

Mechanistic lodging models exist that can predict stem failure from measured stem properties such as yield stress, diameter, and wall thickness (Berry *et al*., 2004), and the empirical literature documents how shading degrades these properties (Luo *et al*., 2022; Zheng *et al*., 2017). However, no model currently predicts stem mechanical properties from the light environment: lodging models require stem properties as inputs rather than simulating them from environmental drivers (e.g., Berry *et al*., 2003). Meanwhile, the crop yield models that underpin current nuclear winter food security assessments — including DSSAT (Blouin *et al*., 2025) and CLM5crop (Xia *et al*., 2022) — do not simulate wheat lodging at all and implicitly assume that plants remain upright through harvest. If low-light lodging occurs at the irradiance levels predicted for temperate regions during the first years of a 150 Tg nuclear winter, these assessments may substantially overestimate temperate wheat production. The same concern may extend to other tall-stemmed cereals, including rice and maize, for which low light has also been shown to reduce lodging resistance (Xue *et al*., 2016; Wu *et al*., 2017), and which together with wheat account for the majority of global staple-crop calories.

## Limitations

CO_2_ concentration in the temperate chambers averaged 552 μmol/mol, well above the current ambient level of approximately 425 μmol/mol. Post-nuclear-war atmospheric CO_2_ would remain near present-day levels, so the elevated concentration was not representative of the target scenario if the war occurred soon. CO_2_ fertilization of C3 crops at this level could increase yields by 10 to 15% relative to ambient (Ainsworth and Long, 2021), making the reported yields slightly optimistic. Similarly, elevated CO_2_ may have partially mitigated lignin depletion under low light, meaning that stem failure could onset earlier or more severely at ambient CO_2_.

The spectral composition of the growth chamber lighting differs from soot-filtered sunlight. No nuclear winter climate simulation has published wavelength-resolved surface irradiance, so the magnitude of any spectral mismatch cannot be quantified. However, at the severe PAR reductions predicted for nuclear winter, total photon flux is likely to dominate over spectral quality as a determinant of yield.

The 16 h photoperiod used in the tropical chamber, while providing a daily light integral equivalent to the predicted equatorial irradiance, does not replicate the approximately 12 h photoperiod of equatorial regions. Photoperiod independently influences flowering time in wheat, and the longer day length may have accelerated development in the tropical chamber, potentially shortening the grain-fill period and affecting yield.

### Implications and future directions

To our knowledge, this experiment provides the first empirical yield data for staple crops under the temperature and light conditions predicted for nuclear winter. Published assessments of food security in nuclear winter have relied on crop modelling alone (Ozdogan *et al*., 2013; Jägermeyr *et al*., 2020; Xia *et al*., 2022; Blouin *et al*., 2025). The present study begins to provide experimental data, though as a controlled environment experiment this is only a first step toward field-scale validation.

Despite the severity of the 150 Tg scenario, the tropical results suggest that meaningful crop production would be possible in equatorial regions. All three crops produced viable yields: wheat yielded approximately 60% of the current global average, while unreplicated canola and potato observations indicated yields near global averages.

In contrast, the temperate results compound an already dire outlook. Under nuclear winter irradiance, wheat stems failed structurally. Although hand-harvesting recovered 0.62 to 2.81 t/ha of grain from the matted canopy (Table 1), mechanical harvest of severely lodged wheat would recover substantially less (Berry *et al*., 2004). Even in the high-light control chamber, where lodging was not observed, winter wheat yielded only 0.86 to 1.35 t/ha (roughly one-quarter to one-third of the current global average of 3.6 t/ha). The Australia site simulated in the temperate chambers was not chosen arbitrarily. Crop models had identified Australia as one of the more promising temperate locations for nuclear winter agriculture (Xia *et al*., 2022; Blouin *et al*., 2025). For the majority of cropland in North America, Europe, and Central and East Asia, the same models predict near-complete loss of production. If wheat cannot be reliably grown even at a site selected for its relative promise, the prospects for temperate cereal production under a 150 Tg nuclear winter are bleak. Canola, however, yielded 2–3 t/ha in unreplicated pots (near current global averages) even under low-light temperate conditions and showed no stem failure, providing preliminary evidence that some crops may tolerate conditions that severely compromise wheat.

These results have direct implications for the crop models that underpin nuclear winter food security assessments. All such assessments (Xia *et al*., 2022; Blouin *et al*., 2025) rely on process-based models that have never been validated under the extreme low-light conditions of a nuclear winter. In particular, the self-weight stem failure observed in temperate low-light wheat represents a failure mode absent from most crop yield models, which implicitly assume that plants remain upright through harvest. Crop model frameworks should treat wheat yield predictions with caution when daily light integrals during stem elongation fall below ∼10 mol m^-2^ d^-1^ (stem failure was not observed at 11.5 mol m^-2^ d^-1^ in the tropical chamber but occurred after approximately 3.5 months below 7 mol m^-2^ d^-1^ in the temperate chamber), though the exact threshold for stem failure remains to be determined. Calibrating crop models for the nuclear winter regime is a necessary next step, but will require substantially more experimental data than this single study can provide.

The challenges of growing crops under nuclear winter conditions documented here underscore that conventional agriculture alone may be insufficient to prevent famine under nuclear winter. A growing body of work has identified a portfolio of complementary food supply interventions that could help close the caloric gap (García Martínez *et al*., 2025), including leaf protein concentrate from legume biomass (García Martínez *et al*., 2026), cellulosic sugar from biomass (Throup *et al*. 2022, Siva and Anderson 2024), greenhouses (Alvarado *et al*., 2020), duckweed farming (Femeena *et al*., 2023; Femeena and Brennan, 2025), seaweed farming (Jehn *et al*., 2024; Hinge *et al*., 2025), wild edible plants (Winstead and Jacobson, 2022), and industrial food production without agriculture (García Martínez, Behr, and Denkenberger, 2024). These interventions could substantially improve food security outcomes under a nuclear winter scenario (Pham *et al*. 2022; Rivers *et al*. 2024).

Several directions for future work emerge from these results. The crop and cultivar space tested here was limited. Additional staple crops, as well as additional wheat, potato, and canola varieties can be tested, ideally with larger sample sizes to move beyond exploratory observations. Testing other cereals (e.g., rye, barley, oats, rice, maize, millet, sorghum) for low-light lodging is a particular priority. Determining which cereals lodge under nuclear winter irradiance, and at what threshold, would clarify whether low-light lodging is a wheat-specific problem or a systemic vulnerability of cereal agriculture under sun-obscuring catastrophes. Short-season crops such as spinach also deserve attention for locations where nuclear winter would leave only a brief window of viable growing conditions.

Future experiments could also measure stem diameter, wall thickness, and lignin content to distinguish the relative contributions of stem geometry and cell-wall composition to low-light lodging. Quantifying the harvest losses specific to low-light lodging would also be valuable. The hand-harvested yields reported here capture the biological grain production of lodged plants but not the additional losses incurred during mechanical harvest of a flat canopy. Paired hand- and machine-harvested plots of lodged wheat could measure this effect. Refrigerated greenhouses equipped with shade cloth and fans to simulate wind could be used to test whether low-light stem failure occurs under realistic mechanical loading. Agronomic and genetic adaptations warrant investigation as potential mitigations: reduced planting density could increase per-plant light interception, while shorter wheat cultivars may reduce the mechanical lever arm and improve stem stability under low light.

More experimental data of the kind reported here are needed to calibrate crop models for the nuclear winter regime, since process-based models remain the only viable tool for assessing food security at global scale. Gaining this knowledge in advance is essential: if a nuclear winter were to occur, knowing which crops and cultivars are most promising for rapid crop switching could materially improve the prospects for sustaining human civilization through such a catastrophe.

## Acknowledgements

Claude Opus 4.6 and 4.7 (by Anthropic) were used to assist with manuscript preparation. We thank Juan B. García Martínez for useful comments.

## Author contributions

Conceptualization: CA, DD, JRL, and SB; methodology: CA, DD, JRL, and SB; investigation: DD, DRA, RB-Z, and SB; formal analysis: SB; writing — original draft: SB; writing — review and editing: all authors; resources: CA and JRL; funding acquisition: CA, DD, and JRL.

## Conflict of interest

The authors declare no conflict of interest.

## Funding

This research was supported by the Food Resilience in the Face of Catastrophic Global Events grant funded by Coefficient Giving (formerly Open Philanthropy) and by the Alliance to Feed the Earth in Disasters (ALLFED).

## Data availability

All harvest measurements and weekly plant height data are provided in a supplementary Excel file.

## Notes

### Competing Interest Statement

The authors have declared no competing interest.

