## Supplementary Figure 1 for "Crop yields under simulated nuclear winter: a growth chamber experiment"

### Supplementary Materials

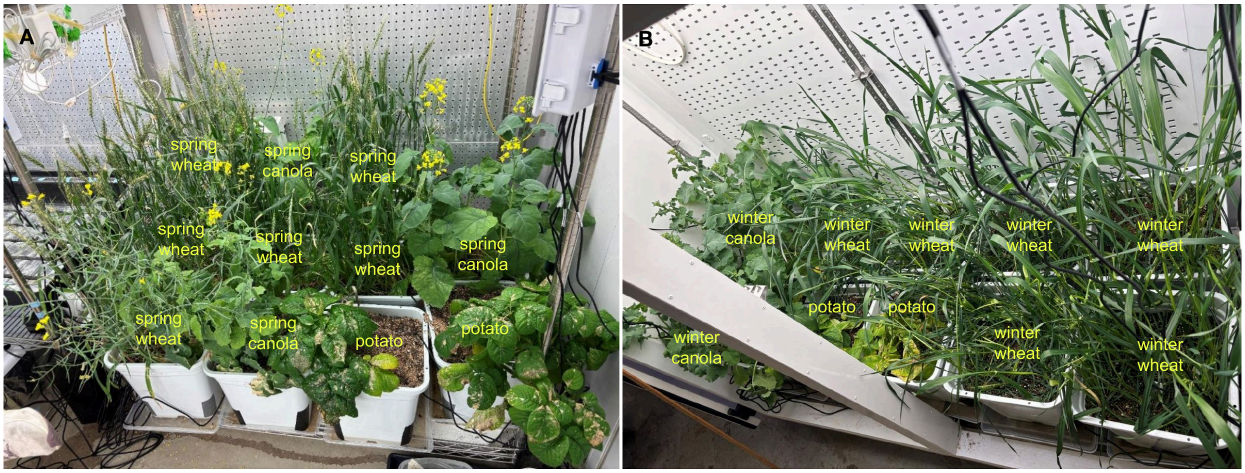


**Supplementary Fig. 1.** Growth chambers used to simulate nuclear winter conditions. (*A*) Tropical chamber (18–20 °C, 200 μmol $m^{-2}$ $s^{-1}$ PAR, 16 h photoperiod) at week 12 after planting, with spring wheat (cv. Glenn), spring canola (cv. NC155 TF), and potato (cv. Red Norland). (*B*) Temperate high-light chamber (variable temperature simulating South Australia under a 150 Tg nuclear winter scenario, constant 360 μmol $m^{-2}$ $s^{-1}$ PAR) at week 26 after planting, with winter wheat (cv. TAM 114), winter canola (cv. Wichita), and potato (cv. Red Norland). The temperate low-light chamber (not shown) had an identical setup. In all chambers, each pot (30.5 × 30.5 × 30.5 cm) contained either nine wheat plants, six canola plants, or one potato plant.
